# Structural basis for EarP-mediated arginine glycosylation of translation elongation factor EF-P

**DOI:** 10.1101/116038

**Authors:** Ralph Krafczyk, Jakub Macošek, Daniel Gast, Swetlana Wunder, Pravin Kumar Ankush Jagtap, Prithiba Mitra, Amit Kumar Jha, Jürgen Rohr, Anja Hoffmann-Röeder, Kirsten Jung, Janosch Hennig, Jürgen Lassak

## Abstract

Glycosylation is a universal strategy to post-translationally modify proteins. The recently discovered arginine rhamnosylation activates the polyproline specific bacterial translation elongation factor EF-P. EF-P is rhamnosylated on arginine 32 by the glycosyltransferase EarP. However, the enzymatic mechanism remains elusive. In the present study, we solved the crystal structure of EarP from *Pseudomonas putida.* The enzyme is composed of two opposing domains with Rossmann-folds, thus constituting a GT-B glycosyltransferase. While TDP-rhamnose is located within a highly conserved pocket of the C-domain, EarP recognizes the EF-P via its KOW-like N-domain. Based on our structural data combined with an *in vitro /in vivo* enzyme characterization, we propose a mechanism of inverting arginine glycosylation. As EarP is essential for pathogenicity in *P. aeruginosa* our study provides the basis for targeted inhibitor design.

## INTRODUCTION

Translation elongation is a non-uniform process and directly depends on the amino acids to be incorporated into the growing polypeptide chain.^1^ Due to its chemical and physical properties, proline delays the peptidyl transfer reaction^2^ and ribosomes can even stall upon translation of distinct diprolyl containing sequence motifs (Figure 1).^3-4^ Such ribosome stalling is alleviated by the eukaryotic and archaeal elongation factor 5A (e/aEF-5A)^5^ and its prokaryotic orthologue elongation factor P (EF-P).^6-12^ The L-shaped EF-P is composed of three β-barrel domains and structurally resembles t-RNA in both size and shape.^13^ EF-P binds to the polyproline-stalled ribosomes between the binding sites of peptidyl-tRNA (P-site) and the exiting tRNA (E-site)^14^ and stimulates peptide bond formation by stabilization of the CCA end of the P-site prolyl-tRNA (Figure 1).^15-16^ A conserved positively charged residue located at the tip of the EF-P KOW-like N-domain is essential for function.^6, 15^ However, for full EF-P activity this residue is post-translationally elongated.^17^ Certain bacteria, including *Escherichia coli* and *Salmonella enterica,* β-lysinylate a conserved lysine K34^EF-P^ by EpmA. This EF-P specific ligase uses β-(R)-lysine as substrate, which is generated by isomerization of α-(S)-lysine employing the activity of the amino mutase EpmB.^18-21^ By contrast, activation of a phylogenetically distinct group of EF-Ps encoded in species such as *Pseudomonas aeruginosa,* or *Neisseria meningitidis*, depends on rhamnosylation of an arginine R32^EF-P^ in the equivalent position.^15, 22-23^ Rhamnosylation is mediated by the recently discovered glycosyltransferase EarP, which inverts the rhamnosyl moiety of the donor nucleotide sugar dTDP-β-L-rhamnose (TDP-Rha) into α-rhamnosyl-arginine when attached to EF-P.^24-25^ Compared to the common and relatively well understood asparagine glycosylation, sugar modifications on the guanidino group of arginine appeared to be rare and almost nothing is known about the molecular mechanism.^26-27^ Beside EF-P arginine-glycosylation, to date there are only two further reported cases: The first one described self ß-glycosylation of sweet corn amylogenin.^28^ In the second case an effector glycosyltransferase termed NleB of enteropathogenic *E. coli* (EPEC) was shown to inactivate human cell death-domain-containing proteins by N-acetylglucosaminylation of arginine and with this being a major pathogenicity determinant during infection.^29-30^ Similarly, a lack of *earP* abolishes pathogenicity of *P. aeruginosa.*^15^ Accordingly, solving the molecular mechanism of arginine rhamnosylation might pave the way to ultimately design and develop targeted inhibitors against EarP.

**Figure 1.**
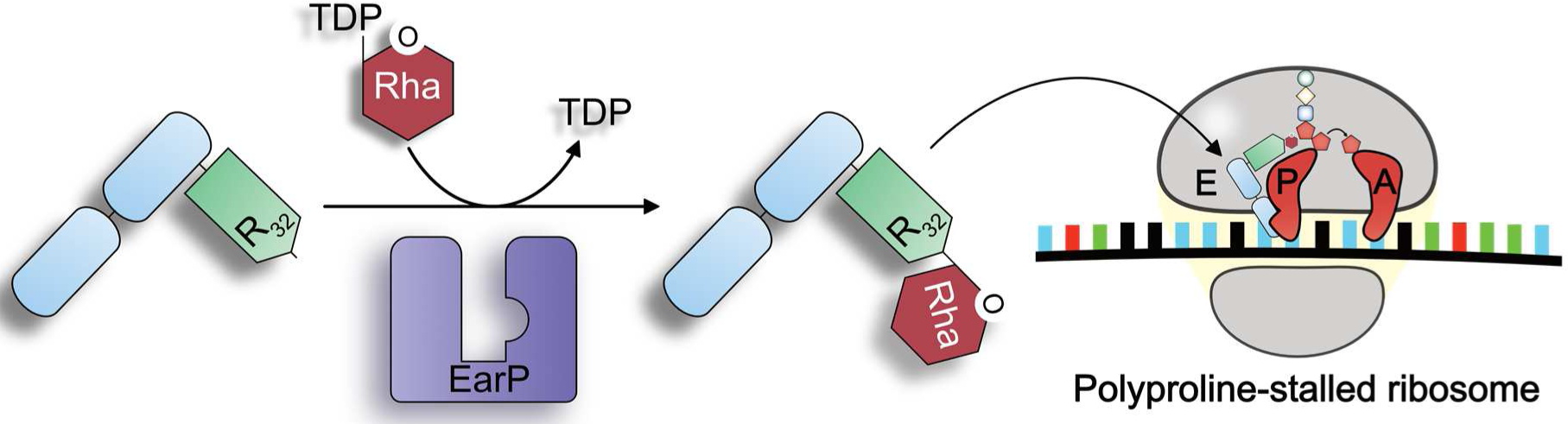
Activation and molecular function of EarP-Arginine type translation elongation factor EF-P. Left: The bacterial translation elongation factor EF-P is composed of two OB-Fold domains (light blue) and one KOW-like N-domain (light green). In about 10% of all bacteria EF-P is post-translationally activated by alpha-glycosylation of a strictly conserved arginine (R32). The glycosylation reaction is catalysed by the EF-P-arginine rhamnosyltransferase EarP using dTDP-β-L-rhamnose (TDP-Rha) as a substrate. Right: Activated EF-P is recruited to polyproline stalled ribosomes and binds between the E and P site. Thereby R32^EF-P^ and the attached rhamnose moiety presumably stabilize the CCA-end of the P-site prolyl-tRNA which in turn stimulates Pro-Pro peptide bond formation and thus alleviates the translational arrest.

Here we present the X-ray crystal structure of EarP from *Pseudomonas putida* KT2440 (EarP_*Ppu*_) bound to its cognate nucleotide-sugar donor substrate TDP-Rha at 2.9 Å resolution (PDB: XXXX). Together with NMR spectroscopy analyses and an *in vitro* / *in vivo* biochemical enzyme characterization, we provide first insights into the mechanism of arginine glycosylation mediated by EarP and thus lay the foundation to understand this yet poorly characterized catalysis.

## RESULTS

Despite low sequence conservation most nucleotide sugar dependent (Leloir-type) glycosyltransferases adopt one of two major folding patterns, GT-A or GT-B.^26^ However, so far there is no available information on the structure and folding properties of EarP. We used SWISS MODEL^31^, Phyre^2 32^ and the I-TASSER server^33-35^ for protein structure and function predictions to generate fold recognition models of EarP from *Pseudomonas putida* (Figure S1). These predictions suggested the UDP-N-acetylglucosamine dependent glycosyltransferases MurG from *E. coli* (MurG_*Eco*_)^36^ and OGT from *Xanthomonas campestris* (OGT_*xca*_)^37^ as structural orthologues. Accordingly, EarP_*Ppu*_ adopts a clamp-like structure with two opposing Rossmann-like domains that are separated by an inter-domain cleft (Figure S1) and with this the protein is presumably a GT-B type glycosyltransferase.^26^

### Structure of *Pseudomonas putida* EarP

We were able to subsequently confirm the GT-B fold by having solved the crystal structure of EarP_*Ppu*_ at 2.9 Å resolution (Figure 2A). Indeed, the EarP_*Ppu*_ C-domain - being very well resolved - includes the residues 184-360 and follows the Rossmann-fold topology with five β-strands (β8-β12) and seven α-helices (α7-α13, Figure 2B). On the other hand, the N-domain (aa 1-153 and 361-377) could only be resolved partly. In the modelled structure, the N-domain features a central β-sheet of seven β-strands (β1-β7), surrounded by five α-helices (α1-α4 and α14) (Figure S1, Figure 2B). In the crystal structure, only β-strands β1, β2, and β3 as well as α-helix α1 and α14 are resolved (Figure 2A). Although, very weak electron density for likely other regions could be noticed, it was not possible to trace the protein sequence due to low resolution and lack of connectivity with the rest of the protein molecule. This weak and discontinuous electron density in several parts of the N-domain suggests that these parts are mobile and could provide an explanation for a higher than usual R-free (37.4%) value at this resolution. In addition, the mobility of the N-domain is further supported by higher average B-factors for this domain compared to the C-domain (61 Å^2^ vs 46 Å^2^; see Figure S2 for B-factors mapped onto the protein structure). However, we were able to confirm the presence of the predicted remaining strands and helices and thus the validity of the model and crystal structure by NMR secondary chemical shifts (Figure 2C). A prerequisite for this analysis is the backbone chemical shift assignment by triple resonance NMR experiments. The relatively large size of EarP_*Ppu*_ with 43 kDa exceeds the sensitivity limitations of NMR demanding for deuteration in order to decrease cross-relaxation effects and to decrease the signal linewidth. Nonetheless, coupled with TROSY-based experiments we were able to assign 62 % of the EarP_*Ppu*_ backbone.

The two domains are interconnected by a bipartite helix (α5, α6) comprising aa 157-176. This linker region together with an unstructured segment that positions α14 in the vicinity of the N-terminus defines the floor of the cleft that separates the domains.

**Figure 2.**
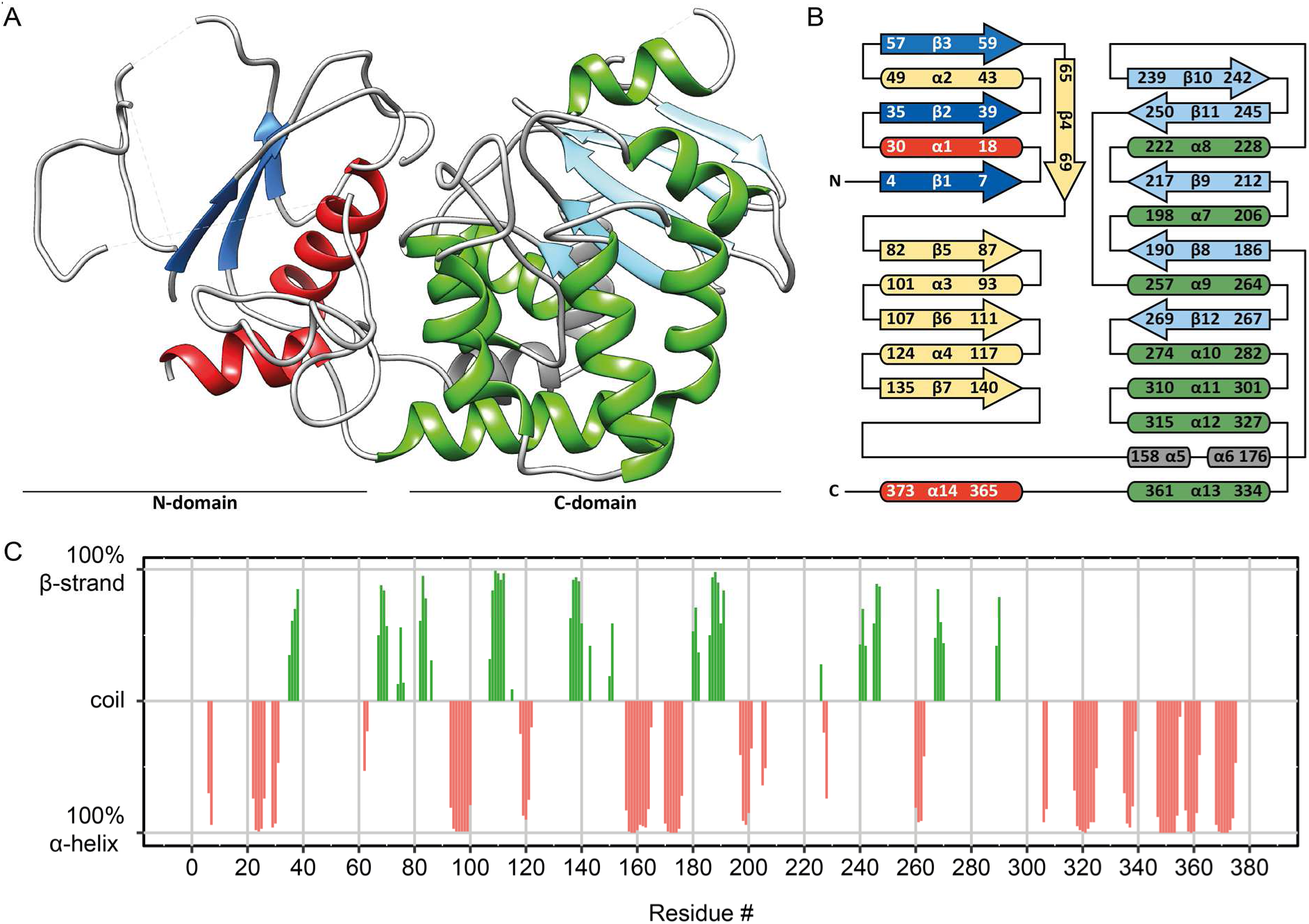
EarP folding pattern and topology. **(A)** Ribbon representation of the 2.9 Å crystal structure of EarP_*Ppu*_. Illustration was generated with UCSF Chimera.^38^ **(B)** Topology diagram of EarP based on the EarP_*Ppu*_ crystal structure, NMR analysis and predictions by MINNOU.^39^ Secondary structure elements are shown, with α-helices in red and green for the N- and C-domains, respectively, and β-strands correspondingly in blue and cyan. The bipartite helix of the linker domain is coloured in grey. Helices and β-strands not resolved in the crystal structure are coloured in yellow. **(C)** Secondary structure of EarP. The secondary structure of individual amino acids is indicated as propensity to either form an α-helix (red) or β-strand (green). The amino acids with a propensity to adopt random coil or lacking information about secondary structure were assigned zero values in the plot. The propensity values were obtained from C_α_, C_β_, NH and H chemical shifts using TALOS+^40^.

### Analysis of the TDP-β-L-rhamnose binding site in the EarP C-domain

In Leloir-type GT-B glycosyltransferases the nucleotide-sugar binding site is canonically located in the protein C-domain.^41^ Similarly, the TDP-Rha in the EarP_*Ppu*_ crystal structure is located in an aromatic pocket that is composed of F191^EarP^, F252^EarP^ and F258^EarP^, surrounding the dTDP moiety, and Y193^EarP^ and Y291^EarP^ on the rhamnose side of the ligand (Figure 3A, Figure S3). The aromatic ring of F252^EarP^ establishes π-stacking with the thymine nucleobase and the ring of F258^EarP^ stacks on the ribose. F191^EarP^ forms a lid on top of the thymine-ribose part. The hydroxyl group at C3’ of the rhamnose is in hydrogen bonding distance with the hydroxyl group of Y193^EarP^ and the aromatic pocket is closed on the rhamnose side by Y291^EarP^. R271^EarP^ forms a hydrogen bond via its guanidino amide bond with the hydroxyl group at C2’ of rhamnose. The binding is further strengthened by side chain interactions of Q255^EarP^, D274^EarP^ and S275^EarP^, forming a hydrogen bond network with the pyrophosphate group.

**Figure 3.**
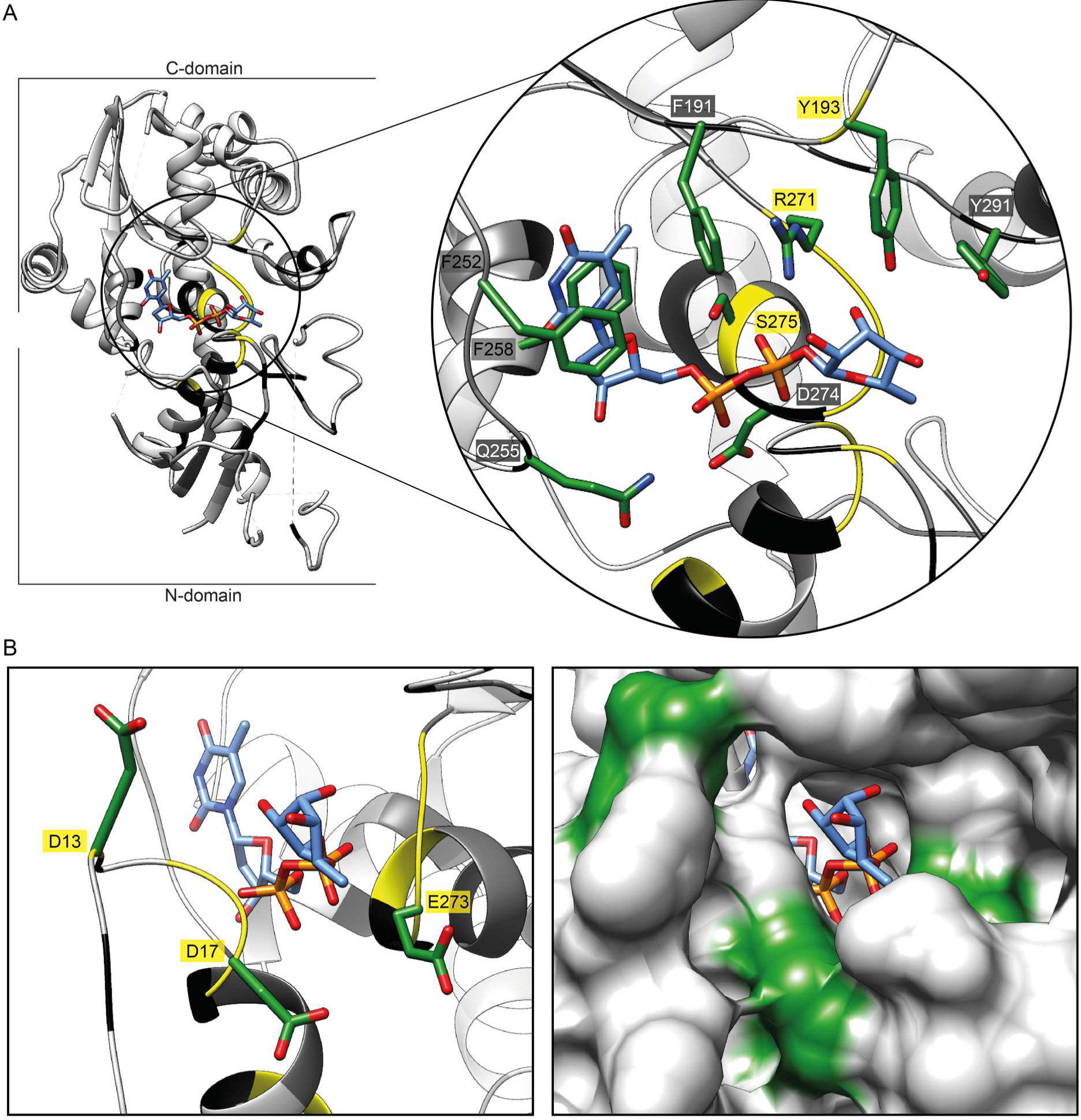
The EarP TDP-β-L-rhamnose binding site. **(A)** Left: Three-dimensional structure of EarP_*Ppu*_ in ribbon representation. The TDP-Rha binding pocket in the C-domain is circled in black. Right: Zoom into the nucleotide-sugar binding pocket with bound TDP-Rha. Important residues for TDP-Rha positioning are depicted as green sticks and labelled with single letter-code identifiers. **(B)** Left: Ribbon representation of the nucleotide-sugar binding pocket with stick representation of the bound TDP-Rha (blue) as well as the three invariant residues D13, D17 and E273 (green) that are presumably involved in catalysis. Right: Surface representation of the nucleotide-sugar binding pocket with stick representation of the bound TDP-Rha (blue), illustrating the closed state of the aromatic pocket in the bound state. D13, D17 and E273 are coloured in green. Backbone residues are colour coded in the ribbon representations according to their degree of conservation: yellow / 100%, black ≥ 95%, dark grey ≥90%, light grey ≥50% and white <50% identical residues in all analysed EarP orthologues. Electron density for TDP-Rha bound to EarP is shown in Figure S3. All illustrations were generated with UCSF Chimera.^38^

To validate the amino acids involved in coordinating the donor substrate in solution, we performed NMR structural analyses including titrations with TDP-Rha. Upon titration with TDP-Rha, clear chemical shift perturbations could be observed, confirming that TDP-Rha binds to EarP at expected residues (Figure 7).

In parallel, small-angle X-ray scattering (SAXS) of free EarP_*Ppu*_ and bound to TDP-Rha has been performed (Figure S4). The overall shape of the molecule could be validated to be the same in solution. Furthermore, there are no large conformational changes (> 10 Å) or movements of both Rossmann fold domains with respect to each other upon binding of TDP-Rha as the scattering density does not change compared to the free state.

Database mining identified 432 EarP homologs representing about 10% of sequenced bacteria (Supplemental Dataset S1).^15^ Phylogenetically EarP originated presumably in the β-proteobacteria subdivision and was horizontally transferred into the γ-proteobacterial orders of *Pseudomonadales, Aeromonadales* and *Alteromonadales.* It can also be found in certain *Fusobacteria, Planctomycetes* and *Spirochetes*.^15^ In order to identify conserved amino acids we used Clustal Omega^42^ and generated a multiple sequence alignment (Figure 4A). We found 49 residues with a sequence conservation of ≥ 95%. Mapping of these residues onto the crystal structure, revealed an accumulation at or near the inter-domain cleft (Figure 4B, C) including the binding pocket for the nucleotide sugar donor substrate (Figure 3, Figure S3).

**Figure 4.**
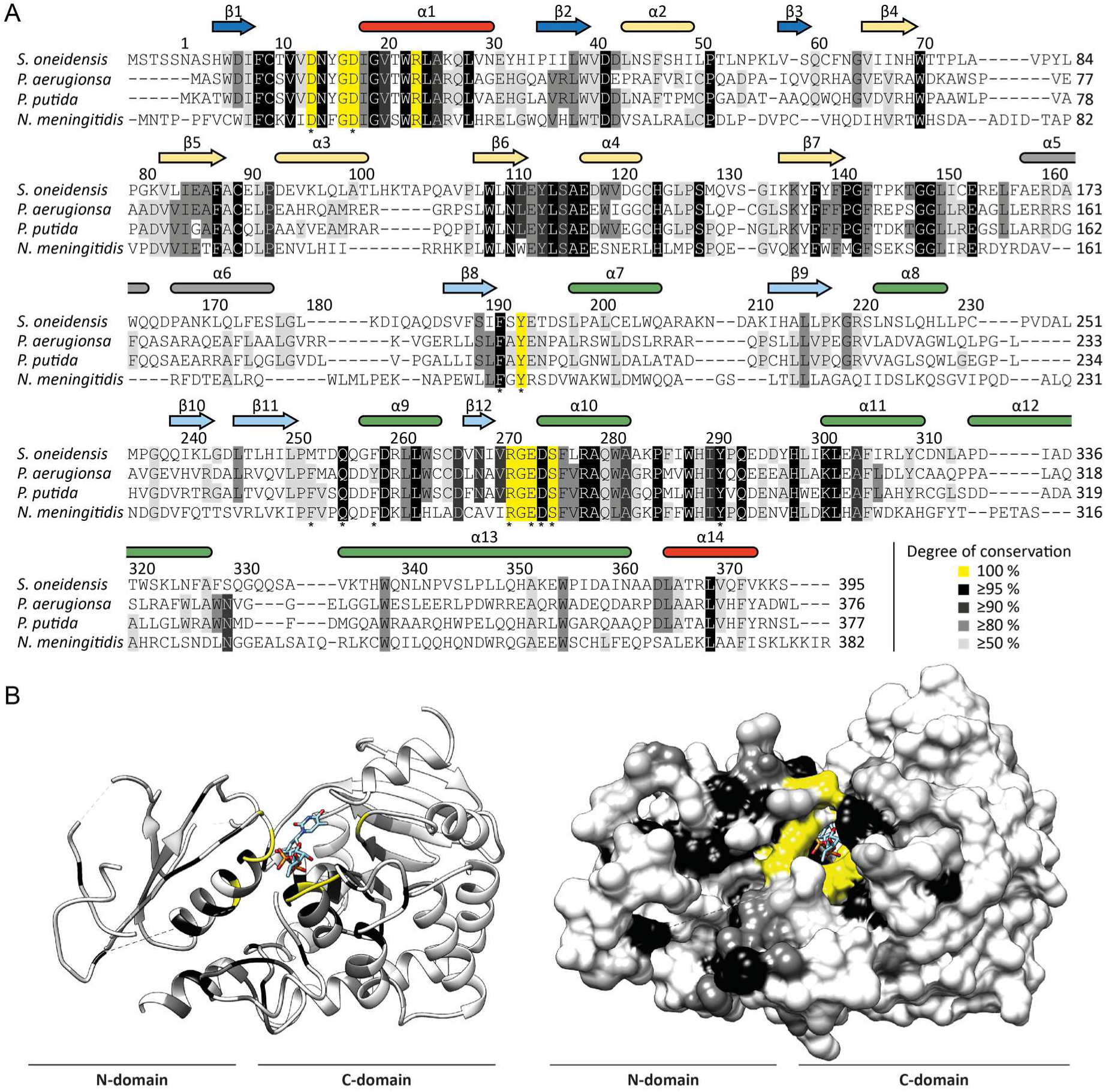
Evolutionary conservation of amino acids in EarP homologs. **(A)** Multiple sequence alignment of EarP from *Shewanella oneidensis, P. aeruginosa*, *P. putida* and *N. meningitides* as a selection from the alignment of 432 protein sequences that were collected from the NCBI database (Supplemental Dataset S1). The multiple sequence alignment was generated using Clustal Omega^43^. Secondary structure elements of EarP are shown with the same colour code as in Figure 2. Amino acids selected for mutational analysis are indicated by asterisks. **(B)** The EarP_*Ppu*_ crystal structure was coloured according the degree of conservation of the respective aa. Ribbon (left) and surface (right) representation of the EarP_*Ppu*_ crystal structure. Residues colour code: yellow 100%, black ≥95%, dark grey ≥90%, light grey ≥50% and white <50% identical residues in all analysed EarP orthologues. Illustrations were generated with UCSF Chimera.^38^

Together with the structural information, this enabled a rational mutational analysis in which selected EarP_*Ppu*_ residues of the TDP-Rha binding site were exchanged by alanine (Figure 4A). This included F191^EarP^, Y193^EarP^, F252^EarP^, F258^EarP^ and Y291^EarP^ forming the aromatic pocket as well as Q255^EarP^, R271^EarP^, D274^EarP^ and S275^EarP^ being involved in hydrogen bond networking. The corresponding EarP_*Ppu*_ substitutions variants were subjected to an *in vivo* functionality analysis (Figure 5A, B).

**Figure 5.**
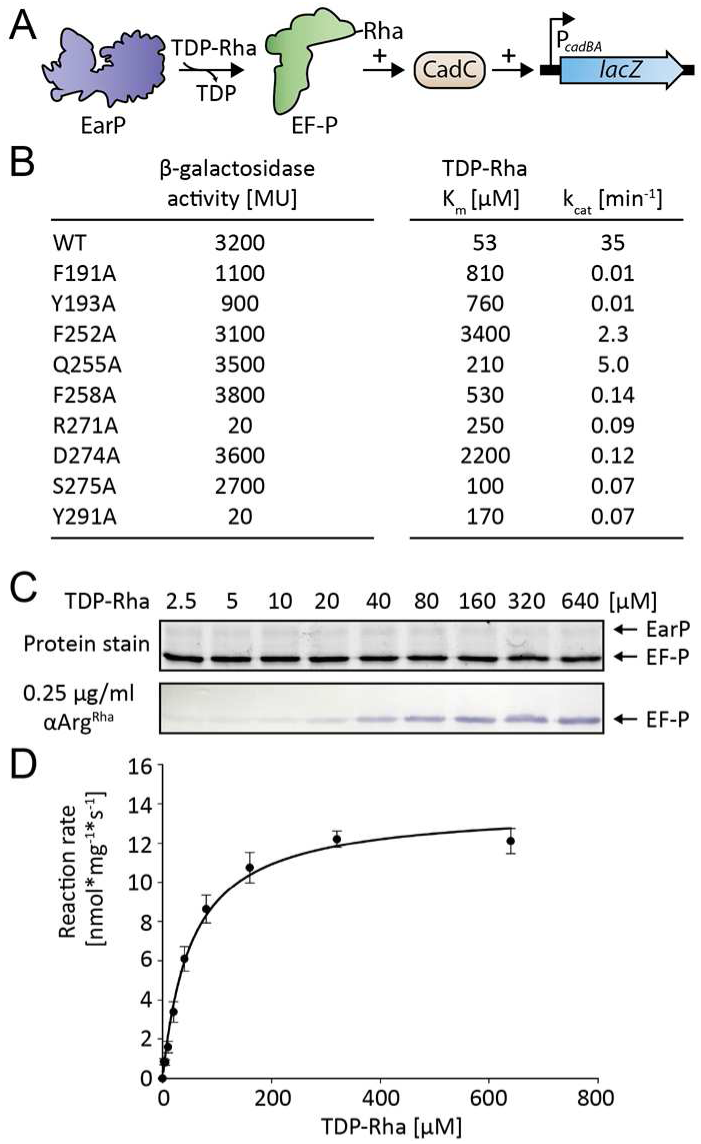
Analysis of kinetic parameters and *in vivo* activity of EarP_*Ppu*_ variants. **(A)** Molecular principle of the *in vivo* EF-P_*Ppu*_ functionality assay. This assay is based on the lysine decarboxylase acid stress response of *E. coli* the CadABC module. At low pH and the concomitant presence of lysine the transcriptional activator CadC activates the promoter of its two downstream genes *P_cadBA_* and with this induces expression of *lacZ* in an *E. coli* MG1655 *P_cadBA_::lacZ* strain. Proper translation of CadC is dependent on the presence of EF-P_*Ppu*_ and its corresponding modification system EarP_*Ppu*_ and thus β-galactosidase activity is a direct read out for EF-P and EarP functionality, respectively. **(B)** I*n vivo* activities and kinetic parameters of tested single amino acid exchange variants of EarP_*Ppu*_. Left: *In vivo* EarP_*Ppu*_ activities were determined by measuring the β-galactosidase activities of an *E. coli* MG1655 P*_cadBA_::lacZ* Δ*efp* heterologously expressing *efp_Ppu_* together with *earP_Ppu_* wildtype or mutants from o/n cultures in LB pH 5.8. Background corrected mean values of three independent measurements are shown. Standard deviations were determined from three independent experiments to be ≦10%; Right: K_m_ and k_cat_ of wildtype EarP_*Ppu*_ (WT^EarP^) and single amino acid substitution variants. Standard errors were determined by SigmaPlot to be <20% **(C)** Top: TCE protein stain^45^ of a representative SDS-gel used for determination of kinetic parameters. Fixed amounts of EF-P_*Ppu*_ (0.25 μM) and WT^EarP^ (0.1 μM) were incubated with varying concentrations of TDP-Rha for 20 s and subjected to SDS-PAGE. Bottom: Detection of rhamnosylated EF-P_*Ppu*_. EF-P_*Ppu*_ was visualized after Western Blotting using 0.25 μg/ml αArg^Rha^ **(D)** TDP-Rha saturation curve of WT^EarP^. Band intensities of (C) were quantified using ImageJ.^46^ Reaction rates were calculated as mean from four independent measurements. Standard deviations are shown as error bars for each concentration.

Previously we could show that the heterologous expression of *efp* and *earP* from *S*. *oneidensis* in *E. coli* can fully complement for a lack of EF-P^15^ with respect to the activation of the lysine dependent acid stress response by the transcriptional activator CadC.^6^ Similarly, coproduction of wildtype EF-P*Ppu* and wildtype EarP*Ppu* (WT^EarP^) can restore wildtype β-galactosidase activity in an *E. coli P_cadBA_::lacZ* Δ*efp* strain (Figure S5). Out of the nine tested EarP_*Ppu*_ substitution variants we measured reduced β- galactosidase activities for F191A^EarP^, Y193A^EarP^, R271A^EarP^, S275A^EarP^, Y291A^EarP^ (Figure 5B). Thereby, R271A^EarP^ and Y291A^EarP^ failed to induce β-galactosidase expression at all.

In parallel the enzymatic activity of EarP*Ppu* was investigated *in vitro* employing an αArg^Rha^ antibody. The antibody was raised against a chemically synthesized glycopeptide antigen (SGR^Rha^NAAIVK) and specifically detects arginine rhamnosylation (see material and method section) (Figure S6). This in turn enabled the quantification of rhamnosylation levels of EF-P_*Ppu*_ over time by Western Blot analysis (Figure 5B, C, D). In a first step K_m_ and k_cat_ of WT^EarP^ were determined to be 53 μM and 35 min^−1^, respectively (Figure 5B, D).

We wondered whether this Km makes physiologically sense and therefore analysed the cellular TDP-Rha levels in *P. putida, P. aeruginosa* and *E. coli,* which were 3.5 mM, 2.0 mM and 4.0 mM, respectively (see material & method section, Figure S7). In good accordance with our measurements in *Lactococcus lactis,* the physiological TDP-Rha concentration was previously determined to be as high as 1 mM.^44^ Thus, within a bacterial cell the donor substrate reaches saturating concentrations according to the WT^EarP^ K_m_ measurements.

Next, K_m_ and K_cat_ of EarP_*Ppu*_ substitution variants were determined and compared to those of the wildtype protein. Strikingly, all mutations affected enzymatic activity (Figure 5B, Figure S8). Depending on the substituted residue the Km increased up to 60fold for F252A^EarP^ (Km=3.4 mM). Conversely, k_cat_ decreased up to 35 times when measuring the kinetics of F191A^EarP^ and Y193A^EarP^, respectively.

To exclude that decreased enzyme activity was due to fold disruption, selected EarP_*Ppu*_ variants (F191A^EarP^, Y193A^EarP^, R271A^EarP^, D274A^EarP^ and Y291A^EarP^) were analysed by NMR ^1^H-^15^N-HSQC experiments (Figure S9). All tested substitution variants showed no structural alterations to the wildtype protein, except for D274A^EarP^. Accordingly, misfolding contributes to the altered enzymatic activity of this protein variant *in vitro.*

Altogether, the *in vivo* and *in vitro* measurements of EarP_*Ppu*_ enzymatic activity complement and validate the structural analysis of the TDP-Rha binding pocket.

### The KOW-like EF-P N-domain is sufficient for EarP mediated rhamnosylation

To test which part of EF-P is involved in the interaction with EarP, NMR chemical shift perturbation experiments were performed by comparing ^1^H-^15^N-HSQC between unbound EF-P_*ppu*_ and EarP_*Ppu*_-bound EF-P_*Ppu*_ (Figure 6A). Triple resonance experiments of EF-P_*Ppu*_ enabled backbone assignment with a sequence coverage of 97 %. Missing assignments are for residues S123, R133, N140, V164, D175 and G185. The assignment enabled also secondary structure determination from secondary chemical shifts and confirmed the validity of the EF-P model for *P. putida,* based on the crystal structure of *P. aeruginosa* EF-P^47^ (Figure S10). The titration experiment showed clear chemical shift perturbations in the N-terminal acceptor domain of EF-P_*Ppu*_ (Figure 6B). However, R32^EFP^ and residues surrounding the rhamnosylation site (e.g. S30^EF-P^ G31^EF-P^ R32^EF-P^ N33^EF-P^) are severely line broadened beyond detection. Therefore, chemical shift perturbation values cannot be determined for these and vicinal residues. This line broadening is an indication that they are bound by EarP_*Ppu*_ and thus have rotational correlation times expected for a complex of that size. Several residues located in the S1-like OB domain are also slightly affected. However, this is not necessarily due to direct contacts with EarP_*Ppu*_ but could also be propagating effects. Therefore, we also investigated *in vivo* and *in vitro* rhamnosylation of truncated EF-P_*Ppu*_ variants comprising either amino acids 1-128 and 1-65, respectively (Figure 6D). Both truncations were readily rhamnosylated by EarP_*Ppu*_ further corroborating that EF-P contact sites are predominantly located in the KOW-like N-domain.

**Figure 6.**
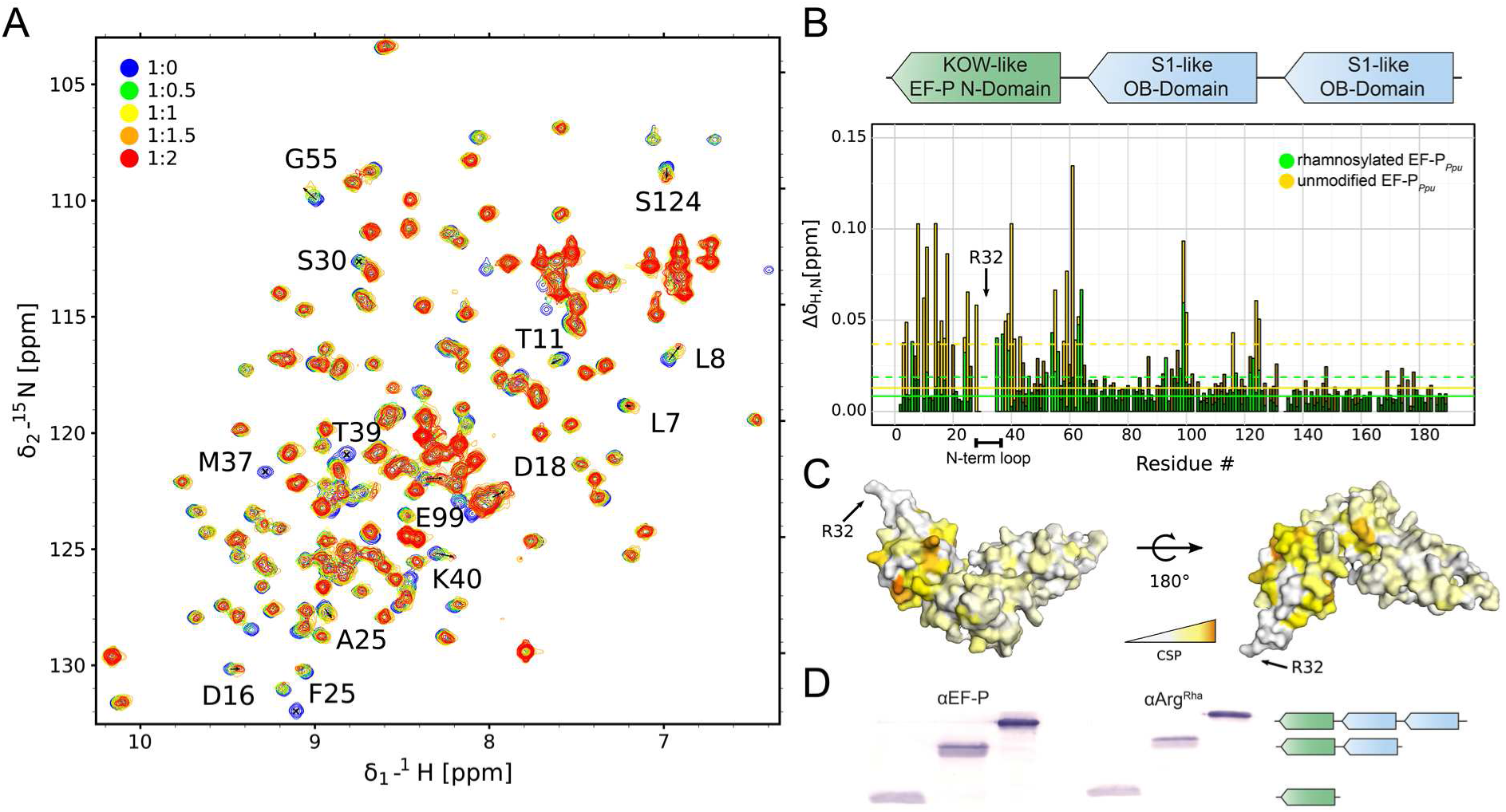
Interaction of EF-P_*Ppu*_ with EarP_*Ppu*_. **(A)** NMR titration of unmodified EF-P_*Ppu*_ titrated by EarP_*Ppu*_. Overlay of ^1^H-^15^N HSQC spectra of EF-P recorded at different titration steps. EF-P was titrated to 1:2 EF-P_*Ppu*_:EarP_*Ppu*_ molar ratio. Colour coding for respective titration steps is indicated in the upper left corner. Examples of peaks with high chemical shift perturbations (CSPs) or severe line-broadening are shown by labels indicating the assignment of given peak. **(B)** Top) Domain structure of EF-P. EF-P consists of three β-barrel domains. The KOW-like EF-P N-domain harbors the rhamnosylation target R32^EF-P^. Bottom) CSPs of EF-P_*Ppu*_ titrated by EarP_*Ppu*_ derived from A). Unmodified and rhamnosylated EF-P_*Ppu*_ were titrated by EarP_*Ppu*_ to 1:2 EF-P_*Ppu*_:EarP_*Ppu*_ molar ratio. To analyse the interaction, CSPs were calculated as described in the methods section and plotted against residue numbers. Colour coding is indicated in the upper right corner. Full lines indicate median CSP, dashed lines indicate median CSP plus standard deviation and residues with CSPs higher than median plus standard deviation are shown in a brighter shade of respective colours. The N-terminal loop containing rhamnosylation target R32^EF-P^ is indicated below the plot. **(C)** CSPs of unmodified EF-P_*Ppu*_ titrated by EarP_*Ppu*_ plotted on the model of EF-P from *P. aeruginosa^47^* (PDB ID: 3OYY) using a white-orange gradient, where white represents the weakest CSP and orange indicates the strongest CSP. The position of R32^EF-P^ is indicated. **(D)** Rhamnosylation experiments using full length EF-P_Ppu_ and C-terminally truncated variants (EF-P_Ppu_ 1-128, EF-P_Ppu_ 1-65). Rhamnosylation of purified protein was detected using 0.25 μg/ml aArg^Rha^. The domain structure of the respective protein variants is indicated as in (B).

In addition, we compared NMR interactions between EarP_*Ppu*_ and unmodified EF-P_*Ppu*_ or rhamnosylated EF-P_*Ppu*_, respectively. This experiment clearly showed that chemical shift perturbations for unmodified EF-P are stronger than for rhamnosylated EF-P (Figure 6B). Thus, EarP releases EF-P after rhamnosylation due to decreased affinity, while unmodified EF-P binds with higher affinity to enable efficient post-translational modification.

### D13, D17 and E273 are involved in catalysis to activate the acceptor substrate EF-P

We and others previously showed that EarP inverts the anomeric configuration on the sugar moiety from TDP-β-L-rhamnose to α-rhamnosyl arginine.^24-25^ Reportedly, inverting glycosyltransferases employ a direct displacement S_N_2-like reaction.^48^ The molecular basis for inverted N-linked glycosylation was elucidated for the oligosaccharyl transferase PglB.^49^ Here the catalytic site features three acidic side chains.^27^ Analogously to PglB, three negatively charged residues - aspartates D13^EarP^, D17^EarP^ and glutamate E273 ^EarP^ - were identified as candidates to catalyze the glycosylation reaction (Figure 2B). All three residues are strictly conserved in all EarP orthologues (Figure 4A, Supplemental Dataset S1). Moreover, D17^EarP^ as well as E273 ^EarP^ are located in proximity to the putative active center for (Figure 3b). Consequently, we constructed the corresponding alanine substitution variants D13A^EarP^ D17A^EarP^ and E273A^EarP^ and investigated the enzymatic activity *in vitro.* In line with the idea that these residues might be involved in catalysis, EF-P rhamnosylation could not be detected even after 8h of incubation and accordingly these EarP variants are inactive (Figure S11). To further confirm our hypothesis, D13A^EarP^, D17A^EarP^ and E273A^EarP^ were investigated on activation of EF-P_*Ppu*_ *in vivo.* Expectedly, coproduction of D13A^EarP^, D17A^EarP^ and E273A^EarP^ with EF-P_*Ppu*_ phenocopies Δ*efp* with respect to *P_cadBA_* activation (Figure 7A; Figure S5) and thus further corroborates our findings from the *in vitro* analysis.

**Figure 7.**
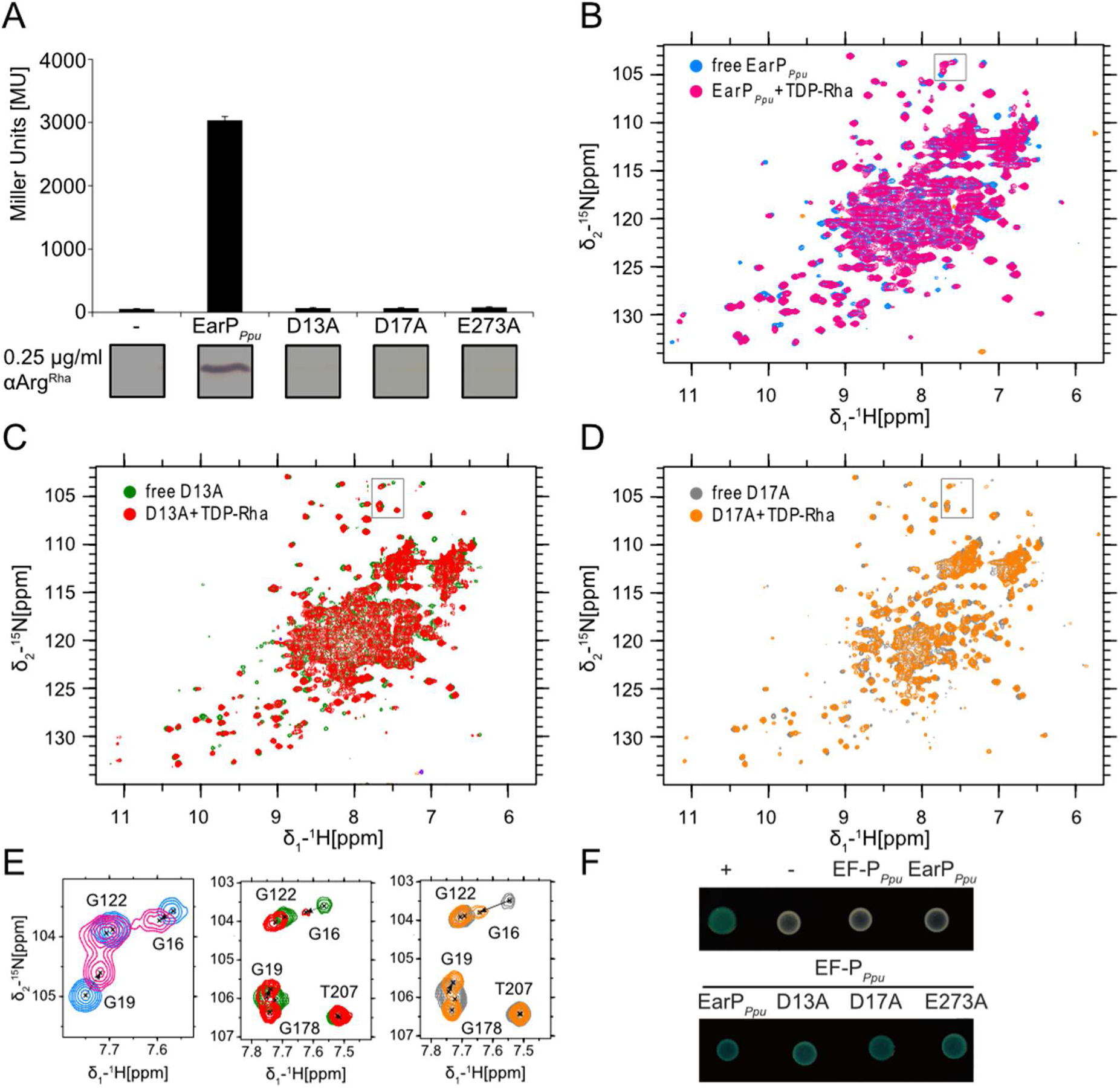
Interaction of TDP-Rha with EarP. (A) **(A)** Top: Analysis of in vivo activity of WT^EarP^, D13A^EarP^, D17A^EarP^ and E273A^EarP^. *In vivo* Ear_*PPpu*_ activities were determined by measuring the β-galactosidase activities of an *E. coli* MG1655 PcadBA::*lacZ* Δ*efp* heterologously expressing *efp^Ppu^* together with *ear_PPpu_* wildtype or mutants from o/n cultures in LB pH 5.8. Mean values of three independent measurements are shown. Standard deviations from three independent experiments were determined to be ≤10% Bottom: Western Blot analysis of o/n cultures of *E. coli* MG1655 P_cadBA_::*lacZ* Δ*efp* heterologously expressing efpPpu together with ear*_PPpu_* wildtype or mutants. Rhamnosylated EF-P^Ppu^ (EF-P^Rha^) was detected by using 0.25 μg/ml αArg^Rha^ **(B)** Overlay of ^1^H ^15^N 301 HSQC spectra of free wild-type EarP and EarP bound to TDP-Rha. **(C)** Overlay of ^1^H ^15^N HSQC spectra of free and TDP-Rha-302 bound D13A EarP variant. (D) Overlay of ^1^H ^15^N HSQC spectra of the free and TDP-Rha-bound D17A EarP variant. The colour coding is indicated in the upper left corner of each spectrum. The titrations are described in details in the material and methods section. **(E)** Zoom-in into the overlaid spectra shown in panels (A), (B) and (C). Position of the zoom-in is indicated by a black frame in the respective original overlay. Peak assignments are shown. The movement of G16 and G19 upon TDP-Rha titration is indicated by dashed arrows **(F)** Bacterial two-hybrid analysis of protein-protein interactions between WT^EarP^, D13A^EarP^, D17A^EarP^ and E273A^EarP^ and the protein acceptor EF-P_Ppu_ in *E. coli* BTH101. The blue colour of colonies results from cleavage of X-Gal by β-galactosidase and indicates protein-protein interaction between co-expressed hybrids.

To exclude misfolding being causative for the non-functional EarP_*Ppu*_ substitution variants, ^15^N-HSQC experiments were performed on D13A^EarP^ and D17A^EarP^, both part of the highly flexible N-domain (Figure 3b). The spectra show no structural alterations compared to WT^EarP^ (Figure 7B and 7C). Additionally, D13A^EarP^ and D17A^EarP^ were titrated with TDP-Rha and are indistinguishable from WT^EarP^ perturbations. Therefore, it can be concluded that donor substrate binding is unaffected. Interestingly, although D13^EarP^ and D17^EarP^ resonances could not be assigned, other residues in close proximity (G16^EarP^ and G19^EarP^) exhibit strong perturbations not only in WT^EarP^ but also in D13A^EarP^ and D17A^EarP^ upon TDP-Rha binding, despite not forming direct ligand contacts. This confirms not only that these mutations do not affect donor substrate binding, but that this region of EarP_*Ppu*_ is also structurally altered upon binding, possibly preparing for efficient EF-P binding (Figure 7D). In parallel, a bacterial two hybrid analysis^50^ was set up to investigate interactions between EF-P_*Ppu*_ and WT^EarP^ as well as D13A^EarP^, D17A^EarP^ and E273A^EarP^. Therefore, fusions were generated with two complementary fragments, T25 and T18, encoding segments of the catalytic domain of the *Bordetella pertussis* adenylate cyclase CyaA. If EF-P*Ppu* and WT^EarP^ do interact, then CyaA is reconstituted, which in turn allows induction of the *lac* promoter and results in *lacZ* expression. Accordingly, the interaction can be visualized by a blue colour of the colony on X-Gal containing plates. Indeed, such blue colonies were not only found when co-producing EF-P_*Ppu*_ with WT^EarP^ but also with the derivatives D13A^EarP^, D17A^EarP^ and E273A^EarP^ (Figure 7G). In comparison; when producing either WT^EarP^ or EF-P_*Ppu*_ together with the corresponding CyaA fragment solely, then the colony remains white (Figure 7G), excluding unspecific interactions.

Thus, altogether these results strongly indicate that D13A^EarP^, D17A^EarP^ and E273A^EarP^ are directly or indirectly involved in activation of the acceptor guanidine group of R32^EF-P^.

## DISCUSSION

Activation of the proline-specific translation elongation factors EF-P and IF-5A is usually achieved by post-translational elongations of the ε-amino group of a conserved lysine.^18-21, 49-50^ The resultant noncanonical amino acids – β-lysinyl-hydroxylysine, hypusine and 5-amino-pentanolyl-lysine – appear to be chemically and structurally analogous. We recently showed that in a subset of bacteria a so far unappreciated form of post-translational modification plays an important role in the activation of EF-P. Here, instead of lysine the guanidine group of a conserved arginine is modified with a rhamnose moiety by a glycosyltransferase termed EarP.^15^ This type of modification does not only contrast with the other known EF-P/IF-5A activation strategies but is also one of only two reported cases of enzyme mediated arginine glycosylation. In the canonical N-linked glycosylation, the sugar is attached to the amide nitrogen of an asparagine in an N-X-S/T consensus sequence (X is any amino acid except for a proline).^48, 53^ Unlike, the effector glycosyltransferase NleB of enteropathogenic *E. coli* N-acetyl-glucosaminylates (GlcNac) modifies specifically the arginines R117 and R235 in the death-domain-containing proteins FADD and TRADD, respectively.^29-30^ This in turn antagonizes apoptosis of infected cells thereby blocking a major antimicrobial host response. Notably, EarP neither shows sequential nor structural homologies to NleB and thus the arginine glycosylation of death-domains and EF-P, respectively, are examples of convergent evolution. Accordingly, one can assume that the molecular mechanisms of the glycosyl transfer reactions differ. In 2016 Wong Fok Lung and co-workers mutated *nleB* and identified certain residues in NleB either interfering with FADD binding or preventing GlcNacylation.^54^ Among the non-functional NleB protein variants they confirmed the importance of two invariant aspartate residues D221 and D223.^30^ A catalytic Asp-X-Asp motif is featured by various GT-A glycosyltransferases. Here, the two negatively charged aspartate side chains coordinate a divalent cation that acts as a Lewis acid to facilitate the nucleotide leaving group departure. Negatively charged amino acids also play important catalytic roles in inverting GT-B glycosyltransferases.^48^ In the case of the metal-independent fucosyltransferase FucT^55^ for example, the side chain carboxyl groups of D13 and E95 could work as base catalysts.^48^ Also the activation of the acceptor amide nitrogen by the lipid donor utilizing bacterial oligosaccharyltransferase PglB depends on the two negatively charged amino acids D56 and E319. These residues abolish the conjugation of the nitrogen electrons and allow the positioning of a free electron pair for the nucleophilic attack onto the anomeric centre of the donor substrate.^27^, ^49^ Analogously the three invariant negatively charged residues D13^EarP^, D17^EarP^ and E273^EarP^ in the EarP protein family might play a role in activating the R32 guanidino group of EF-P. Presumably, D17^EarP^ and E273^EarP^ - both being in close proximity to each other and the rhamnose moiety of TDP-Rha - form the catalytic dyad (Figure 2A). However, at this stage it cannot be excluded that D13^EarP^ is also directly involved in catalysis after conformational rearrangements of the β1-β2 loop upon EF-P binding.

While activation of the acceptor substrate could be driven by the essential amino acids D13^EarP^, D17^EarP^ and E273^EarP^ the nucleotide sugar donor TDP-Rha is bound in a highly conserved cavity of the protein C-domain. A co-crystal structure of the putative structural EarP analogue MurG_*Eco*_ with its cognate substrate reveals that aromatic amino acid side chains play important roles in UDP binding.^56^ Similar interactions where reported for the protein O-fucosyltransferase POFUT1, where F357 is involved in Π-stacking with the respective nucleobase ^57^. Stacking interactions also play a role in EarP where the aromatic side chains of F252^EarP^ and F258 ^EarP^ bind the thymine and ribose moiety of TDP-Rha, respectively. By contrast, contacts with the ribose or the phosphate moieties frequently occur via interactions with sidechain amines, hydroxyl groups and backbone amides^36^, ^56-57^. Accordingly, this is also the case for EarP.

In GT-B glycosyltransferases, positively charged amino acids are often involved in facilitating leaving group departure. This is achieved by neutralization of evolving negative charges on the phosphate moiety during the glycosyl transfer reaction as described e.g. for R261 of MurG_*Eco*_.^36^ Notably, *earP_Ppu_* encodes an invariant R271^EarP^ in the equivalent position and a substitution to R271A^EarP^ strongly impairs protein function, altogether suggesting a similar role in product stabilization.

In GT-B glycosyltransferases the two Rossmann folds can be generally divided into one donor and one acceptor substrate binding domain.^41^ Similar to other glycosyltransferases the nucleotide sugar is bound by the protein C-domain of EarP. Accordingly, it is worth assuming important binding sites for EF-P in the protein N-domain. Conversely, EF-P presumably contacts EarP by amino acids that are in close proximity to the glycosylation site R32^EF-P^. In agreement with this hypothesis the EF-P β-lysine ligase EpmA for example recognizes EF-P via identity elements in a region located around the *E. coli* EF-P modification site K34.^19-20, 58^ Along the same line the deoxyhypusine synthase (DHS) can efficiently modify a human eIF-5A fragment comprising only the first 90 amino acids of the protein.^59^ Similarly, we could show that the KOW-like N-terminal domain of EF-P (Figure 6D) is sufficient to be glycosylated by EarP being congruent with the NMR titrations of EF-P with EarP (Figure 6A, B). Upon titration with EarP the chemical shift perturbations observed were - with a few exceptions - restricted to the first 65 residues.

Taken together we propose a three-step model for the rhamnosylation of EF-P by its cognate modification system EarP (Figure 8). In the ground state both the nucleotide sugar binding site in the C-domain and the putative acceptor binding site in the N-domain are unoccupied. In the donor bound state TDP-Rha is coordinated within a highly conserved cavity in the protein C-Domain including an aromatic pocket that surrounds the thymine ring (Figure 3). Previous studies showed that, binding of the donor substrate induces structural alterations in both the N- and C-domains of glycosyltransferases.^41^, ^60-61^ In MurG these rearrangements include rotation of F244 that stacks over the nucleobase to cap the donor binding pocket.^36^ Notably in the crystal structure of EarP a phenylalanine F252 is in the equivalent position indicating that this capping interaction is conserved (Figure 3C).^56^ As mentioned above the structural alterations might not be restricted to the protein C-Domain but can include an up to 20° rotation of the N-Domain to prone the glycosyltransferase for catalysis.^61^ This idea is also conceivable for EarP as the protein N-domain, which presumably binds EF-P, remains highly flexible even in the crystal. Such a movement could relocate the putative catalytic amino acids D13, D17 and E273 thereby forming the actual active site. In this catalytic state (Figure 8C) the R32^EF-P^ guanidino group might be activated especially by interactions with the negatively charged side chains of D17^EarP^ and E273^EarP^ by a mechanism analogous to the one that was reported for the oligosaccharyltransferase PglB.^49^ In the EF-P rhamnosylation reaction, R271^EarP^ might stabilize the nucleotide product and thereby facilitates leaving group departure. Upon successful inverting glycosyl transfer from TDP-Rha to R32^EF-P^ presumably by a single S_N_2 displacement reaction, the products are released from the active site of EarP in turn reverting back to the unbound ground state.

**Figure 8.**
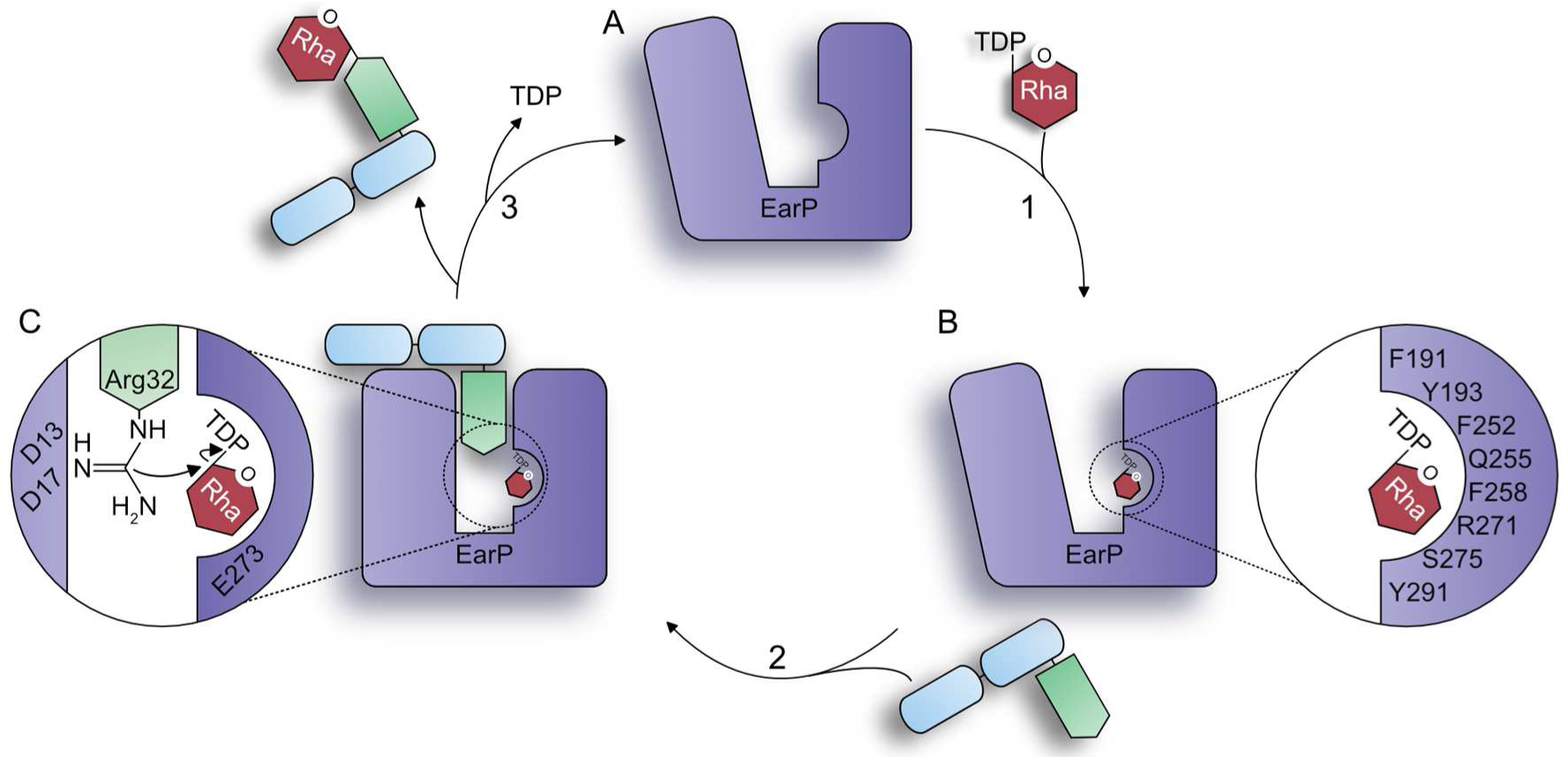
Proposed glycosylation mechanism of EarP. **A)** Ground state; both the donor and acceptor binding site are unoccupied. **B)** Donor-bound state; TDP-Rha is bound and oriented within the binding pocket in the protein C-domain. **C)** Catalytic state; The putative catalytic amino acids D13, D17 and E273 facilitate the nucleophilic attack onto the anomeric centre of TDP-Rha by activating the Arg32 guanidino group. The EarP N-domain is stabilized upon binding of EF-P. Binding and dissociation events indicated by arrows: 1, TDP-Rha binding; 2, EF-P^-^ binding, 3, EF-P^Rha^ and TDP dissociation.

Altogether, our structural and biochemical investigation of EarP provides first insights into the molecular mechanism of arginine glycosylation and with this improves our general understanding in N-linked glycosylation. Additionally, our research might open up new avenues for the development of antimicrobial drugs in order to fight e.g. *P. aeruginosa* infections.

## MATERIALS AND METHODS

### Bacterial strains and growth conditions

Strains and plasmids used in this study are listed in Table S1. *P. putida* and *E. coli* were routinely grown in Lysogeny broth according to the Miller modification^62^ at 30 °C (for *P. putida)* and 37 °C (for *E. coli),* unless indicated otherwise. When required, media were solidified by using 1.5% (wt/vol) agar. If necessary, media were supplemented with 50 μg/ml chloramphenicol, 100 μg/ml kanamycin sulphate, and/or 100 μg/ml ampicillin sodium salt. For promoter induction from plasmid *PBAD* containing plasmids, L-arabinose was added to a final concentration of 0.2% (wt/vol) in liquid medium. For promoter induction from plasmids comprising the *lac* operator sequences, Isopropyl β-D-1-thiogalactopyranoside (IPTG) (Sigma Aldrich) was added to a final concentration of 1 mM in liquid medium.

#### Molecular biology methods

Enzymes and kits were used according to the manufacturer’s directions. Genomic DNA was obtained according to the protocol of Pospiech and Neumann^63^ and plasmid DNA was isolated using a Hi-Yield plasmid Mini Kit (Suedlabor). DNA fragments were purified from agarose gels by employing a High-Yield PCR clean up and gel extraction kit (Suedlabor). Restriction endonucleases were purchased from New England Biolabs (NEB). Sequence amplifications by PCR were performed utilizing the Q5 high-fidelity DNA polymerase (NEB) or the OneTaq DNA polymerase (NEB), respectively. Mutations were introduced into the *earP* gene by overlap extension PCR.^62^ All constructs were analysed by Sanger Sequencing (LMU Sequencing Service). Standard methods were performed according to Sambrook & Russel.^64^

#### β-Galactosidase activity assay

Cells expressing *lacZ* under control of the *cadBA* promoter were grown in buffered LB (pH 5.8) overnight (o/n) and harvested by centrifugation. β-galactosidase activities were determined as described in^66^ in biological triplicates and are given in Miller units (MU).^67^ The significance of the results was determined by applying two-sided students t-test and stating a result as significantly different if p < 0.05.

#### Bacterial two-hybrid analysis

Protein-protein interactions were detected using the Bacterial Adenylate Cyclase Two-Hybrid System Kit (Euromedex) according to the product manuals. Chemically competent^68^ *E. coli* BTH101 cells were cotransformed with pUT18C*-efp_Ppu_* and/or the respective pKT25 variants (pKT25-earP, pKT25-D13A, pKT25-D17A, pKT25-E273A) and plated on LB screening medium containing 40 μg/ml 5-bromo-4-chloro-3-indolyl-β-D-galactopyranoside (X-gal), 0.5 mM IPTG as well as kanamycin sulphate and ampicillin sodium salt. Transformants containing pUT18- *zip* and *pKT25-zip* were used as positive controls. Transformants carrying pUT18C and pKT25 vector backbones were used as negative controls.

Bacteria expressing interacting protein hybrids exhibit a blue phenotype on screening plates due to functional complementation of the CyaA fragments (T18 and T25). After 48 h of incubation at 30 °C plates containing around >100 colonies were evaluated. Representative colonies were transferred to liquid LB culture containing kanamycin sulphate and ampicillin sodium salt and incubated o/n at 30 °C. Subsequently 2 μl of the ON culture were spotted on LB X-Gal/IPTG plates. Pictures were taken after 48 h of cultivation at 30 °C (Figure 7).

#### Protein purification

C-terminal His6-tagged EarP_*Ppu*_ variants (pBAD33-*ear*P_*Ppu*_) were overproduced in *E. coli* LMG194 by addition of 0.2% arabinose to exponentially growing cells and subsequent cultivation at 18°C o/n. N-terminal His6-tagged EarP (pACYC-DUET-*ear*P_*Ppu*_.) and His_6_-SUMO-tagged EF-P_*Ppu*_ (pET-SUMO-*efp_Ppu_*) were overproduced in *E. coli* BL21 (DE3) by addition of 1 mM IPTG to exponentially growing cells. Subsequently, cells were incubated at 18 °C overnight. Rhamnosylated EF-P_*Ppu*_ (EF-P^Rha^) was produced by co-overproduction with His6-tagged EarP_*Ppu*_. Cells were lysed by sonication and His6-tagged proteins were purified using Ni-NTA (Qiagen) according to manufactures instructions. The His6-SUMO-tag was removed by incubation with 1 u/mg His_6_-*ulp*1^69^ overnight. Subsequently Tag free EF-P_*Ppu*_ was collected from the flow through after metal chelate affinity chromatography. For biochemical analyses cells were cultivated in LB. For use in NMR spectroscopy cells were grown in M9 minimal medium.^62^ If necessary ^15^N labelled nitrogen (^15^NH4Cl) and ^13^C labelled glucose were used. For NMR backbone assignment of EarP_*Ppu*_ additionally 99.8 % pure heavy water D_2_O (Sigma-Aldrich) was used instead of H_2_O in growth medium to allow partial deuteration of the protein in order to reduce cross-relaxation effects and increase signal-to-noise. For the production of selenomethylated EarP_*Ppu*_, *E. coli* BL21(DE3) cells expressing N-terminal His6-tagged EarP_*Ppu*_ were cultivated in 1 L M9 minimal medium at 37 °C to an OD600 of 0.6. 100 μg threonine, 100 μg lysine and 50 μg isoleucine were added to feedback inhibit methionine biosynthesis.^70^ Additionally, 50 μg L-(+)-selenomethionine was added 15 min prior induction. Protein production was induced by addition of 1 mM IPTG and cells were incubated at 18 °C overnight. Protein concentrations were determined as described by Bradford.^71^ For biochemical analyses EarP_*Ppu*_ and EF-P_*Ppu*_ were dialyzed against 100 mM NaPi pH 7.6, 5 mM DTT whereas a buffer composed of 100 mM NaPi pH 7.6, 50 mM NaCl and 5 mM DTT was used when the proteins were subjected to NMR analysis.

#### Synthesis of a single rhamnosyl-arginine containing glycopeptide

Moisture and air sensitive reactions were conducted in flame-dried glassware under an argon atmosphere. Commercially available reagents and solvents were used without further purification. CH_2_Cl_2_ was distilled from calcium hydride and THF was distilled from sodium/benzophenone immediately prior to use. DMF was stored under argon in a flask containing 4 Å molecular sieves. Reactions were monitored by TLC with precoated silica gel 60 F254 aluminum plates *(Merck KGaA,* Darmstadt) using UV light and methoxyphenol reagent (100 mL 0.2% ethanolic methoxyphenol solution and 100 mL 2 M ethanolic sulfuric acid) as visualizing agent. Flash chromatography was performed using silica gel (35–70 μm) from *Acros Organics.* Peptide purification by RP-HPLC was performed on a *JASCO* purification system with a UV-Vis detector (model UV-2075Plus) using a *Phenomenex Aeris Peptide* 5u XB-C18 column (250 × 21.2 mm). Analytical RP-HPLC was measured on a *JASCO* system with a *Phenomenex Aeris Peptide* 5u XB-C18 column (250 × 4.6 mm). In all cases, mixtures of water (eluent A) and acetonitrile (eluent B) were used as eluents; if required, 0.1% formic acid (FA) or 0.1% trifluoroacetic acid (TFA) were added. HR-ESI mass spectra were recorded on a *Thermo Finnegan LTQ FT* mass spectrometer or on a *Bruker maxis* equipped with a *Waters Acquity UPLC* using a *Kinetex* C18 column (2.6 μ, 100 A) at 40 °C.

**Scheme 1:**
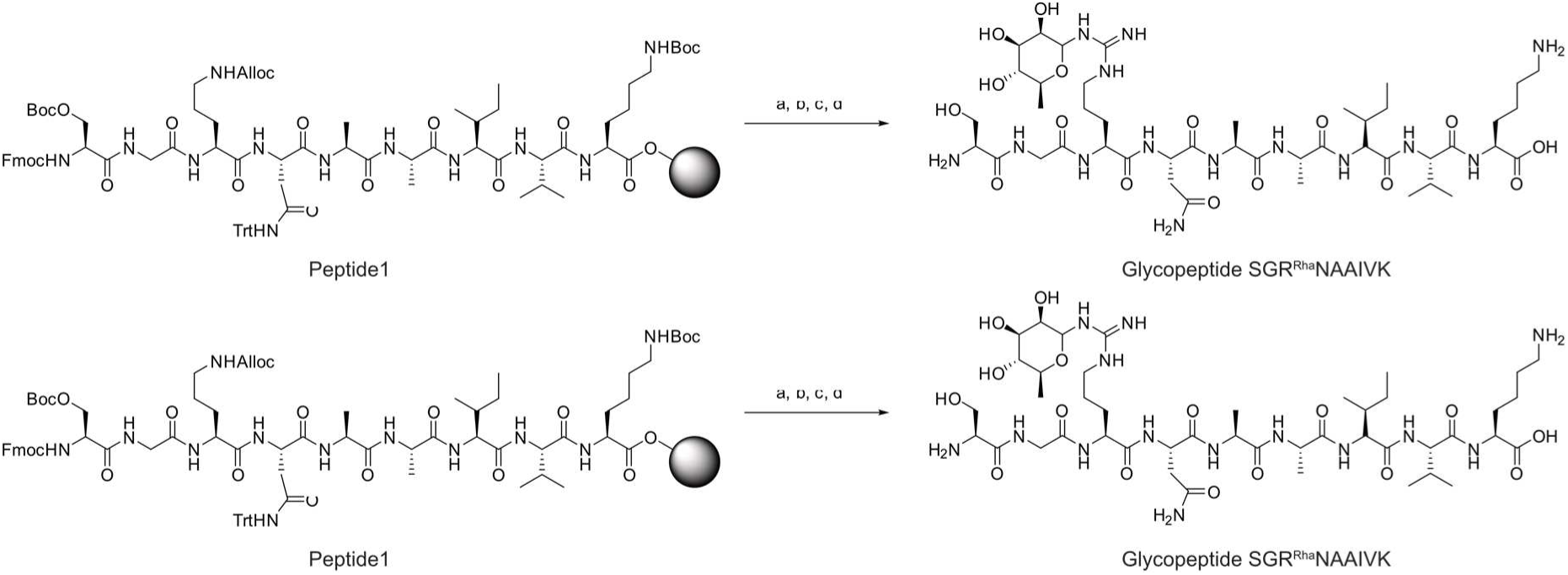
Synthesis of glycopeptide SGR^Rha^NAAIVK; a) SiPhH_3_, Pd(PPh_3_)_4_, CH_2_Cl_2_; b) 1-(*tert*-Butoxycarbonyl)-3-(2,3,4-Tri-*O*- acetyl-6-deoxy-L-mannopyranos-1-yl)-2-ethyl-isothiourea^24^, AgNO_3_, NEt_3_, DMF; c) N_2_H_4_ • H_2_O (5% solution in DMF); d) TFA/H_2_O/phenol/TIPS (88/5/5/2).

Glycopeptide SGR^Rha^NAAIVK was synthesized using a *Liberty Blue™* automated microwave peptide synthesizer followed by *on-resin* glycosylation and deprotection (scheme 1). For construction of peptide 1 0.1 mmol of preloaded H-Lys(Boc)-2-chlorotrityl resin (loading 0.78 mmol/g) were applied. Cleavage of Fmoc protecting group was achieved with 20% piperidine in DMF (75 °C, 35 W, 3 min). Fmoc-protected amino acids (5 eq.) were activated for peptide coupling using 5 eq. of ethyl (hydroxyimino)cyanoacetate (*OxymaPure^®^*), 0.5 eq. of DIPEA and 5 eq. of *N*,*N′*-Diisopropylcarbodiimide. All coupling reactions were conducted at 75 °C and 28 W for 5 min. Removal of the allyloxycarbonylprotecting group and subsequent coupling of the sugar moiety as well as deprotection of the acetyl groups were performed according to established procedures.^24^ Final deprotection gave the desired glycopeptide SGR^Rha^NAAIVK in 39% yield after HPLC purification. The amino acid sequence of the glycopeptide corresponds to the primary structure of the S. *oneidensis* acceptor loop which is highly similar to the consensus sequence of EarP-arginine type EF-Ps.^15^

##### HRMS (ESI+)

Calculated for C_44_H_82_N_14_O_16_ [M+2H]^2+^:m/z = 531.3011; found: 531.3016.

**HPLC** (0.1% TFA, 0 min: 8% B →45 min: 50% B, flow: 1 mL/min): *t_R_*= 9.61 min, *λ =* 204 nm.

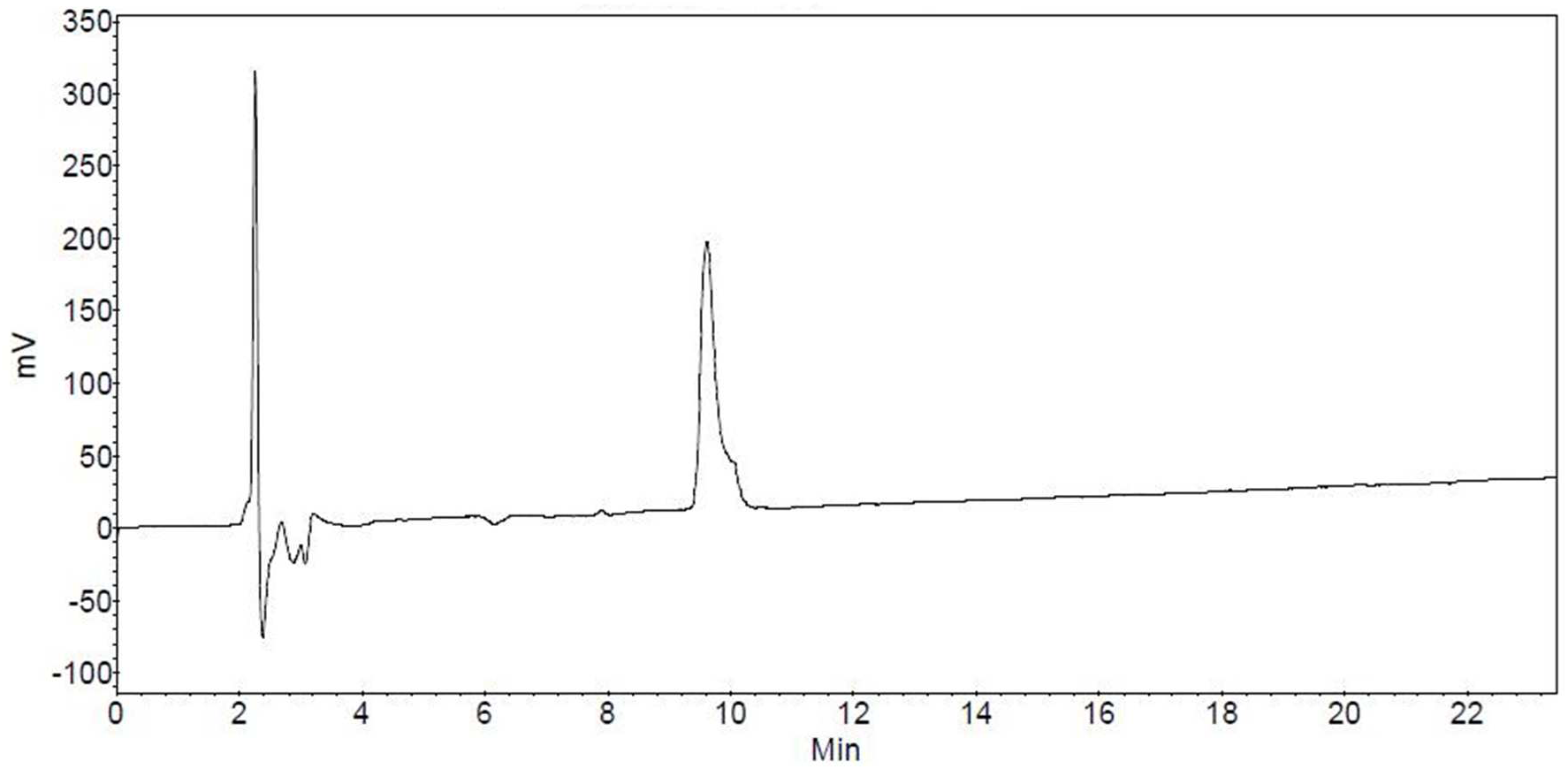

### Antibody generation

Polyclonal antibodies were raised commercially by Eurogentec according to the Rabbit – speedy 28 days (AS-Super-Antigen) program. The mono-rhamnosyl-arginine containing peptide was coupled to BSA according to an internal protocol (AS-PECO-05). Antibodies capable of binding to rhamnosyl-arginine were purified from rabbit sera by affinity chromatography (AS-PURIMED) against the glycopeptide SGR^Rha^NAAIVK. To test the specificity of the purified polyclonal antibodies towards EF-P^Rha^, 1.5 μg of unmodified and 0.5 μg of modified EF-P were transferred to nitrocellulose membrane by Western Blotting. While polyclonal antibodies raised against EF-P from *S. oneidensis* detect both unmodified and modified EF-P_*Ppu*_, our αArg^Rha^ very specifically detect only the modified protein variant (Figure S6).

### SDS-PAGE and Western Blot

Electrophoretic separation of proteins was carried out using SDS-PAGE as described by Lämmli.^72^ Separated proteins were visualized in-gel using 0.5% (v/v) 2-2-2- trichloroethanol^45^ and transferred onto nitrocellulose membrane by vertical Western blotting. Antigens were detected using 0.1 μg/ml Anti-6X His tag^®^ (Abcam), 0.2 μg/ml αEF-P^24^ or 0.25 μg/ml of αArg^Rha^ respectively. Primary antibodies (RABBIT) were targeted by 0.2 μg/ml alkaline phosphatase conjugated Anti-RABBIT IgG (H&L) (GOAT) antibody (Rockland) and target proteins were visualized by addition of substrate solution (50 mM sodium carbonate buffer, pH 9.5, 0.001% (w/v) Nitro-Blue-Tetrazolium, 0,045% (w/v) 5-Bromo-4-chloro-3-indolylphosphate).

### Determination of kinetic parameters

Kinetic parameters were determined by varying TDP-Rha concentrations but keeping concentrations of EarP_*Ppu*_ (0.1 μM) and unmodified EF-P_*Ppu*_ (2.5 μM) constant. A mixture of EarP_Ppu_ and unmodified EF-P_*Ppu*_ was equilibrated to 30°C in 100 mM NaP_i_ pH 7.6. The reaction was started by addition of TDP-Rha and was stopped after 20 seconds of incubation at 30 °C by the addition of one volume 2x Lämmli buffer^72^ and incubation at 95 °C for 5 minutes. Samples were subjected to SDS-PAGE and rhamnosylated EF-P_*Ppu*_ was detected as described above. Band intensities were quantified using ImageJ.^46^ Product formation (nmol*mg^-1^) was calculated relative to fully (*in vivo*) rhamnosylated EF-P_*Ppu*_. K_m_ and K_cat_ values were determined by fitting reaction rates (nmol*mg^-1^*s^-1^) to the Michaelis-Menten equation using SigmaPlot. Time-course experiments conducted at a TDP-Rha concentration of 500 μM show that the rhamnosylation reaction is not saturated after 20 seconds of incubation (Figure S12).

### Fold Recognition

Fold recognition models were generated using the online user interface of PHYRE^2 32, 73^, SWISS-MODEL^74-77^ and the I-TASSER^33-35^ server as instructed on the website. Model structures were selected from the array of results according to best confidence-, qmean- and z-scores respectively. All images of tertiary protein structures in this work were generated using the UCSF Chimera package developed by the Resource for Biocomputing, Visualization, and Informatics at the University of California, San Francisco.^38^ Protein structures were obtained as .pdb files from www.rcsb.org^78^ or the respective modelling platforms mentioned above.

### Determination of intracellular TDP-Rha concentrations

Cells were grown in 1 L LB to an OD_600_ of 0.5 (5*10^8^ cells/ml), harvested by centrifugation and resuspended in 25 ml 100 mM NaP_i_ pH = 7.6 (2*10^10^ cells/ml). After disruption of cells using a Constant Systems Ltd continuous flow cabinet at 1.35 kBar, cell debris were removed by centrifugation and lysates were sterilized by filtration (Steriflip^®^). A mixture of EarP_*Ppu*_ (0.1 μM) and unmodified EF-P_*Ppu*_ (2.5 μM) was equilibrated to 30°C in 10 μl 100 mM NaP_i_ pH = 7.6. The reaction was started by addition of 10 μl lysate from ~2*10^7^ or ~2*10^8^ cells respectively and stopped after 20 seconds of incubation at 30 °C by addition of one volume 2x Lämmli buffer^72^ and incubation at 95 °C for 5 minutes. In parallel, a TDP-Rha calibration series was generated by addition of TDP-Rha at final concentrations ranging from 5 μM to 160 μM including the linear range of the rhamnosylation reaction rate (Figure 4). Samples were subjected to SDS-PAGE and rhamnosylated EF-P_*Ppu*_ was detected as described above. Band intensities were quantified using ImageJ.^46^ TDP-Rha concentrations in samples containing lysate were calculated by dividing the respective relative band intensities by the slope of the corresponding calibration curve (5 μM to 80 μM TDP-Rha). Intracellular TDP-Rha concentrations were calculated from the amount of substance per cell while assuming equal contribution of TDP-Rha across all cells as well as an average cell volume of 3.9 μm^3^ ^79^ for *E. coli* and 2.1 μM^3^ ^80^ for *P. putida* and *P. aeruginosa* respectively.

## NMR spectroscopy

### Backbone assignment of EF-P and EarP

All NMR experiments were performed at 298K on Bruker Avance III spectrometers with a magnetic field strength corresponding to a proton Larmor frequency of 600 MHz equipped with a Bruker TXI cryogenic probehead, 700 MHz equipped with a Bruker room temperature probehead or 800 MHz equipped with a Bruker TXI cryogenic probehead. All datasets were processed using NMRPipe.^76^

Before NMR measurements of ^15^N- and ^13^C labelled EF-P (700 μM) in 100 mM NaP_i_, 50 mM NaCl and 5 mM DTT, pH 7.6, 0.02 % NaN_3_ was added to the sample. Sequential resonance assignment was obtained from 2D ^1^H-^15^N-HSQC, 3D HNCA, CBCACONH, and HNCACB experiments, using constant time during ^13^C evolution.^81^ The assignment process was assisted by CARA (http://cara.nmr.ch) and CcpNmr Analysis^82^ a7 % was obtained. Missing assignment for residues other than prolines are S123, R133, N140, V164, D175 and G185. Secondary chemical shift analysis was performed based on the difference of measured ^13^C_α_ and ^13^C_β_ chemical shifts to random coil chemical shifts of the same nuclei to assign secondary structure to the EF-P sequence (Figure S10) and confirm the validity of the model shown in Figure 6.^83-84^

Due to the size of EarP (43 kDa) backbone resonance assignment was only possible for ^2^H, ^15^N, ^13^C-labelled samples to reduce the number of protons and thus cross-relaxation effects, which also enables efficient acquisition of backbone assignment experiments in TROSY-mode^85^. TROSY-HNCA, - HNCACB, and −CBCACONH experiments^86^, processed by NMRPipe^87^ and analysed using CARA (http://cara.nmr.ch) enabled backbone resonance assignment of 62 % of all assignable residues (excluding prolines).

The NMR titrations were always performed by adding unlabelled interaction partner to ^15^N-labelled protein sample and monitoring the progress of the titration by recording ^1^H-^15^N HSQC. First, ^15^N-labelled 150 μM unmodified EF-P was titrated with unlabelled EarP to 1:2 EF-P:EarP molar ratio with intermediate steps at 1:0, 1:0.5, 1:1 and 1:1.5 EF-P:EarP molar ratio. ^15^N-labelled 41 μM rhamnosylated EF-P was titrated with unlabelled EarP to 1:2 EF-P:EarP molar ratio without any intermediate steps. ^15^N-labelled 540 μM wild-type EarP was titrated with unlabelled TDP-Rha to 1:5 EarP:TDP-Rha molar ratio with intermediate steps at 1:0, 1:0.2, 1:1 and 1:3 molar ratio. ^15^N labelled 186 μM D13A or 209 μM D17A EarP variants were titrated by the addition of TDP-Rha to approximately 1:10 molar ratio with no intermediate steps. To analyse the EF-P:EarP and wildtype EarP:TDP-Rha titration the chemical shift perturbations were calculated according to formula: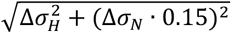, where 0.15 is the weighing factor to account for nitrogen resonances generally spanning a broader frequency range.

To check proper folding of EarP variants ^1^H-^15^N HSQC spectra of ^15^N-labelled EarP variants were recorded for following single amino acid substitutions at following concentrations: 209 μM D13A, 209 μM D17A, 162 μM F191A, 197 μM Y193A, 139 μM D274A, 186 μM R271A and 162 μM Y291A.

## Small-angle X-ray scattering

Thirty microliters of EarP, EarP + TDP-rhamnose, and buffer (with and without TDP-rhamnose) were measured at 20°C at the BioSAXS beamline BM29 at the European Synchrotron Radiation Facility using a 2D Pilatus detector. For each measurement ten frames with 1 s exposure time per frame were recorded for each EarP and buffer sample, using an X-ray wavelength of λ = 0.9919 Å. Measurements were performed in flow mode where samples are pushed through a capillary at a constant flow rate to minimize radiation damage. The protein concentrations measured were 1.0, 2.0, 4.0 and 8.0 mg/ml. TDP-Rha was used in 7:1 excess (ligand protein), respectively. The buffer measurements were subtracted from each protein sample and the low Q range of 1.0 mg/ml was merged with the high Q range of the 8.0 mg/ml sample, using PRIMUS^88^. The merging was done due to the rising scattering density at low Q ranges for the higher concentrated samples, indicative of aggregation. CRYSOL^89^ was used to fit the back-calculated scattering densities from the crystal structure to the experimental data.

## X-ray crystallography

For crystallization N-terminal His_6_-tagged EarP_*P.p*_ expressed as a seleno-methionine derivative was used. The protein was dialysed to 50mM Tris, 100mM NaCl, 1mM DTT, pH 7.6, concentrated to 183 μM and TDP-Rha was added to final concentration of 10 mM. The crystallization condition was 0.2M ammonium acetate, 0.1M bis-tris pH 6.0 and 27% (w/v) PEG 3350. A Full dataset was collected at the ID29 beamline, ESRF, Grenoble at a wavelength of 0.9793111Å (absorption peak for selenium) and 15.05% beam transmission with 0.15° oscillation range, 0.037s exposure time and 2400 frames. The space group was determined to be I4. The dataset was phased using single anomalous dispersion (SAD) by Crank2^90^ automatic pipeline in CCP4^91^ using Afro provided by Pannu, N.S. (unpublished) for FA estimation, Crunch2^92^ for substructure detection and Solomon^93^ for density modification. The initial structure was built in Phenix Autobuild^94^ and was completed by several rounds of manual model building in Coot^95^ and refinement in Phenix^94^.

## ACCESSION CODES

Atomic coordinates of EarP_*Ppu*_ have been deposited in the Protein Data Bank with accession number XXX, while NMR backbone chemical shifts for EarP_*Ppu*_ and EF-P_*Ppu*_ are available at the BMRB (accession numbers: XXXXX and XXXXX).

## ACKNOWLEDGEMENTS

We want to thank Ingrid Weitl for excellent technical assistance. We thank Wolfram Volkwein for fruitful discussions. The SAXS and X-ray diffraction experiments were performed on beamlines BM29 and ID29, respectively, at the European Synchrotron Radiation Facility (ESRF), Grenoble, France.

We are grateful to Local Contacts at the ESRF for providing assistance in using beamlines BM29 and ID29.

## AUTHOR CONTRIBUTION

AHR, SW and DG performed the organic synthesis and NMR analysis of small molecules and wrote the corresponding material and method section of the manuscript. R. K. performed the confirmation of antibody specificity raised against the rhamnosyl arginine comprising peptide. Additionally, RK constructed the EarP_*Ppu*_ and EF-P_*Ppu*_ encoding plasmids and purified all proteins used for biochemical analyses, NMR studies and X-ray crystallography. RK also performed the biochemical *in vivo* / *in vitro* characterization of EarP_*Ppu*_ and determined concentrations of TDP-β-L-rhamnose in *E. coli, P. putida* and *P. aeruginosa.* TDP-β-L-rhamnose was synthesized by JR, PM and AKJ. JH and JM performed and analysed all protein NMR experiments. The crystallisation screen was set up by JM. JH, JM and PKAJ solved the crystal structure of EarP_*Ppu*_. JL, JH and KJ designed the study. The manuscript was written by RK, JM, KJ, JH and JL.

